# EpiTADformer: A Transformer-Based Model for High-Resolution TAD Boundary Detection Using Epigenomic Signal Embeddings

**DOI:** 10.64898/2026.01.20.700691

**Authors:** My Nguyen, Shaojun Tang, Joseph L. McClay, J. Chuck Harrell, Mikhail G. Dozmorov

## Abstract

The human genome is partitioned at different levels of 3D genome organization, with topologically associating domains (TADs) being among the most well-known and biologically important structures. TAD boundary disruption is associated with a wide range of diseases such as cancer, neurological and developmental disorders. Numerous methods have been developed to detect TAD boundaries from chromatin contact maps obtained with Hi-C technology. However, these methods are largely limited by the resolution of Hi-C data, typically 1 Kb to 100 Kb. In contrast, functional DNA loci, collectively referred to as epigenomic data, are profiled at a much higher resolution (100-200 bp for a typical ChIP-seq experiment). To improve the resolution of boundary detection, we hypothesize that the patterns of epigenomic signals associated with regions in proximity to TAD boundaries can serve as embeddings for these genomic regions, defining region similarity. These embeddings, along with their positional relationships, can be effectively modeled using deep learning to achieve more precise boundary prediction. We present EpiTADformer, a transformer-based model that takes as input transcriptional and histone modification signals of neighboring regions centered around TAD boundaries. We demonstrate that EpiTADformer outperforms feedforward neural network, convolutional neural network (CNN), and bidirectional long short-term memory (BiLSTM) network architectures. These results suggest the positional information of epigenomic signals surrounding TAD boundaries provides a strong predictive signal, enabling improved performance of the transformer model. Our findings highlight the potential of epigenomic signals to serve as region embeddings for refining the epigenomic language of TAD domains and 3D genome organization.

## Introduction

The three-dimensional (3D) organization of chromatin plays a fundamental role in key cellular processes such as gene regulation, replication timing, and cell differentiation [1–4]. Hi-C, a genome-wide chromosome conformation capture technique, has enabled the study of chromatin architecture at various levels [5], including A/B compartments, topologically associating domains (TADs), and chromatin loops. Among these findings, TADs—distinct genomic regions containing chromatin with frequent internal interactions—have been linked to the molecular mechanisms of disease [3]. Disruption of TADs can lead to gene dysregulation and contribute to various diseases [3,6]. TADs facilitate precise enhancer-promoter communication, regulate gene expression by spatially organizing regulatory elements, and provide functional insulation between neighboring domains [1,5,7,8]. TAD boundaries—the regions separating TADs—also play a crucial role in gene regulation by limiting the interaction of cis-regulatory elements to their target genes. These boundaries are remarkably stable across cell types and are enriched for CTCF binding, housekeeping genes, evolutionary constraints, and complex-trait heritability, underscoring their functional and evolutionary importance [9].

The study of TAD boundaries and their structures is an active area of research, essential for uncovering the complexities of the human genome and its role in disease mechanisms. The majority of methods for identifying TAD boundaries (e.g., SpectralTAD [10], TADfit [11], RefHiC [12], deepTAD [13], reviewed in Xu et al. 2024 [14]) rely on chromatin contact maps, but their resolution is constrained by Hi-C data, which typically ranges from 1 Kb to 100 Kb [11]. Another group of methods utilize epigenomic data known to be associated with TAD boundaries (e.g., CTCF, RAD21, SMC3, ZNF143 [17,18]) to predict boundaries without the need for Hi-C maps (Hi-ChIP-ML [19], PredTAD [20], reviewed in [21]). Similar to Hi-C-based methods, many epigenomic-based methods have limited resolution (e.g., 10 kb for PredTAD). preciseTAD [22] has emerged as a transfer learning framework for learning TAD boundary association with epigenomic annotations at the Hi-C resolution and predicting boundaries at the base-level resolution. As with other methods, it considered epigenetic signals at each region (base) independently. Methods for high-resolution TAD boundary prediction that consider the continuity of epigenetic signals remain limited [21].

This work presents EpiTADformer, a method for improved TAD boundary prediction utilizing positional information about epigenomic signals (transcription factor binding sites (TFBSs), histone modification marks). EpiTADformer is a transformer encoder-based model that learns both linear and non-linear positional relationships between epigenomic features associated with high-resolution (100 bp) genomic regions (bins). We consider region-specific epigenomic features (e.g., vectors of TFBS signals) as region embeddings capturing similarity among regions. Intuitively, due to continuity of epigenomic signals, two adjacent regions would have similar epigenomic signal profiles, analogous to embeddings of similar words. On the other hand, distant regions, e.g., a region at a boundary and a region at a TAD’s center, would have different epigenomic feature vectors due to different biochemical properties of the genome at those regions, analogous to embeddings of different words. Leveraging this embedding idea, the model is trained on epigenomic signal profiles within windows of regions centered around TAD boundaries (positive examples) vs. randomly selected (negative examples). This approach allows us to improve the location of known TAD boundaries as well as discover candidate novel boundaries enriched in molecular drivers of TAD boundary formation. EpiTADformer can detect TAD boundaries of varying sizes, providing an epigenomic context-informed and high-resolution framework for studying 3D genome organization.

## Results

### Epigenomic embeddings for TAD boundary prediction

Given known associations of TAD boundaries with CTCF, cohesin, H3K27ac, H3K4me3, and other epigenomic features [17,18,23,24], our goal is to use epigenomic signals at and around TAD boundaries to predict TAD boundary locations at high resolution. Epigenomic signals are continuous, with nearby genomic regions typically having similar biochemical properties captured in epigenomic profiles. Therefore, we binned the genome into fixed-size bins (100 bp) and consider a bin-specific epigenomic signal vector as a region embedding vector capturing similarity of nearby regions and dissimilarity of distant regions (we use “bins” and “regions” interchangeably to refer to discrete genomic segments). An analogy can be drawn with word embeddings which capture word similarity and distinguish words with opposite meanings. To illustrate this concept, we selected regions overlapping with known markers of TAD boundaries (CTCF, RAD21, SMC3), combined them with the same number of randomly selected regions, and visualized the t-SNE and UMAP representations of them. As expected, these regions occupied different spaces on the dimensionality reduction plots (Supplementary Figure S1A). Differential analysis of epigenomic signals between positive and negative regions expectedly identified CTCF, SMC3, RAD21 as most significantly different and found other known marks of TAD boundaries, such as YY1, TRIM22, SMARCA5, and others [18]. These results were observed irrespective of chromosome and data transformation (Supplementary Figure S1B,C). These results indicate that epigenomic signal vectors can serve as region embeddings, and their positional relationship may help to improve the detection of TAD boundaries by deep learning models.

The workflow of EpiTADformer is illustrated in Figure 1. The input consists of genomic bins with each bin annotated with epigenomic signal data (e.g., transcription factor binding sites and/or histone modification signals). By default, the genome is processed at the 100 bp bin resolution, a typical resolution for modern chromatin profiling technologies (e.g., ChIP-seq, ATAC-seq). The feature x bin matrix is normalized using z-score transformation by default. Each bin is labeled as boundary or non-boundary based on the boundary points defined by preciseTAD, a machine learning framework shown to outperform other methods for boundary predictions [22]. We defined windows of length *n*_*bins*_ = (2*k*+1), where *k* denotes the number of bins included on each side of the center bin. A window is defined as a boundary if it contains a center boundary bin (positive examples); otherwise it is a non-boundary window (negative examples). All boundary and an equal number of randomly selected non-boundary windows are then combined and randomly shuffled to construct the training dataset (Figure 1A).

To capture the local context of epigenomic signals around TAD boundaries, we adopt a transformer architecture with positional encoding capturing the position of each bin within a window (Figure 1B). The region embeddings and their positional encoding are combined and passed through a transformer encoder block consisting of multi-head self-attention, layer normalization, and feed-forward neural network layers. The output from the transformer encoder is passed through several fully connected layers (MLP, Multi-Layer Perceptron), which further transform the learned representations for the final classification task. At the output layer, predictions for each window are used to define the TAD boundary status of their center bins. Predicted boundary and non-boundary bins that occur in close proximity are then grouped into boundary and non-boundary regions (Figure 1B).

**Figure 1.**
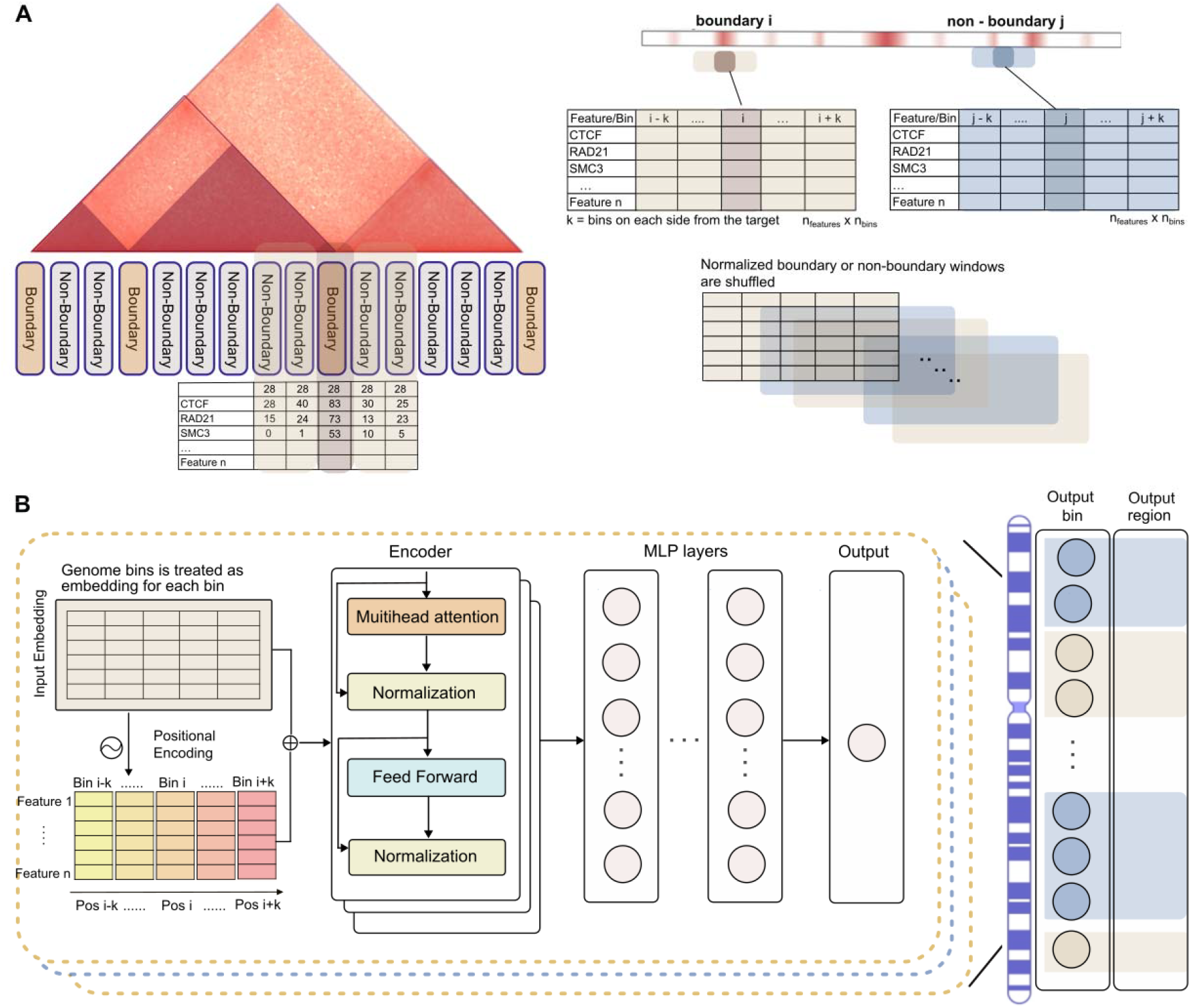
Overview of EpiTADformer. (A) Boundary and non-boundary windows capturing epigenomic signals at and around center bins. Left: The genome is divided into fixed-size bins. For each boundary or non-boundary bin, a fixed-size window (default number of bins in each window is 101) centered on it is extracted. For each window, epigenomic signals (e.g., CTCF, RAD21, SMC3, etc.) across are assembled to form a feature matrix indexed by signal values and bin position ( matrix). Right: Example windows for a boundary (at bin) and a non-boundary region (at bin ) are shown. All normalized boundary and non-boundary matrices are then randomly shuffled to construct the training dataset containing an equal number of positive and negative samples. (B) The epigenomic signal vectors are provided as embeddings for each bin, and positional encoding is added to preserve spatial information of each bin within the window. The encoder applies multi-head attention, normalization, and feed-forward layers to capture both local and long-range dependencies across bins. The encoded output is then passed through multilayer perceptron layers (MLPs) to generate the classification prediction for each bin. Predicted boundary and non-boundary bins that occur in close proximity are further grouped into boundary and non-boundary regions of varying sizes.

**Supplementary Figure S1.**
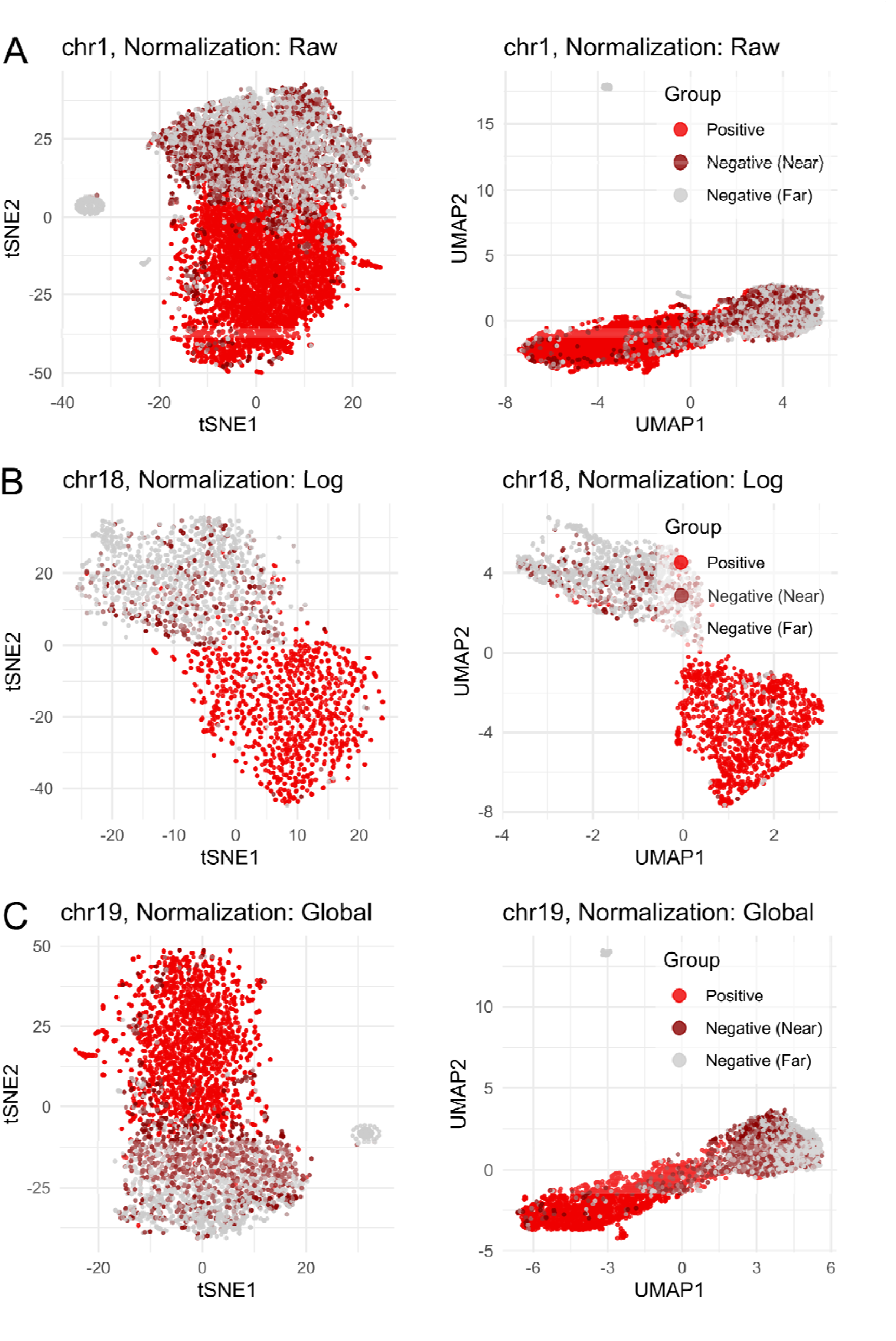
Low-dimensional representation of genomic bin embeddings. t-SNE and UMAP visualization of epigenomic signal vectors for positive (CTCF, RAD21, and SMC3 overlapping), and random regions. The random regions were colored dark red if they were within 50 Kbp of the positive regions (colored red), or gray otherwise. A small number of random regions without any signal formed a separate cluster to demonstrate the importance of signal values in the embedding space. Data for GM12878 cell line, 188 transcription factor binding profiles, 100 bp resolution. (A) chr1 with the largest number of positive regions, raw epigenomic signal data; (B) chr18, least gene dense chromosome, log-transformed data; (C) chr19, most gene dense chromosome, globally normalized data.

### Positional Epigenomic Embeddings Enable Superior EpiTADformer Performance

We aim to demonstrate the ability of epigenomic region embeddings and their positional relationship to predict TAD boundaries. For this, by default, we constructed a z-score–normalized training dataset using windows of 101 bins at 100-bp region size resolution and seven epigenomic signal tracks corresponding to three transcription factors (main transcriptional features - GM12878’s CTCF, RAD21, and SMC3), which have been shown to be strong predictors of TAD boundaries [22]. The training procedures for all models are described in the Methods section.

To evaluate the effect of data preprocessing on the performance of deep learning models, we first compared the performance of baseline models (NN, CNN, Bidirectional RNN) and the Transformer architecture trained on raw, log-transformed, globally normalized epigenomic signal x binned region matrices. We found that models trained on the raw, untransformed data outperformed those trained on log-transformed inputs. We further compared raw versus globally normalized inputs (across bins and features) and observed that most models exhibited improved performance when global normalization was applied (Supplementary Table S1). However, combining global normalization and log-transformation did not lead to a performance increase (Figure 2A). These results suggest the z-score-transformed epigenomic signal range improves model accuracy as compared to raw or log-transformed signal.

We next investigated the effect of window size by varying the number of bins included in each input window. Most models showed improved performance as window size increased, until reaching optimal performance at 101 bins and declining thereafter (Figure 2B). At this optimal window size, the Transformer model achieved the highest performance among all models, with mean F1 score 0.921 as compared with F1 = 0.913 for the NN, 0.914 for the CNN, and 0.918 for the Bidirectional RNN (Supplementary Table S2). However, the Transformer model was also more sensitive to deviations from the optimal window length, with performance degrading more rapidly than the other architectures. This performance degradation is potentially due to larger windows covering additional multimodal signal patterns that may distort the relationship between boundary bins and their nearby features (Figure 1C, Supplementary Table S2). These results highlight the Transformer’s ability to leverage well-defined local context (within 10,100 bp when using windows containing 101 bins) while underscoring the importance of selecting biologically meaningful window sizes.

To assess how bin resolution influences boundary detection, we trained all models using binned signal data at 50 bp, 100 bp, 200 bp, and 400 bp resolutions under the default setting (101 bin windows). Across all bin resolutions, the Transformer model achieved the highest performance, with median F1 scores of 0.88 at 50 bp, 0.92 at 100 bp, 0.92 at 200 bp, and 0.92 at 400 bp (Supplementary Table S3). Model performance increased by approximately 0.1 in median F1 when the resolution was increased from 50 bp to 100 bp, reflecting the benefit of reduced noise and sharper localization of signal peaks (Figure 2D). Beyond 100 bp, performance remained stable, with only minor fluctuations at 200 bp and 400 bp (Figure 2D). These results indicate that the models—particularly the Transformer, which consistently outperformed the others—can effectively learn boundary-associated patterns across a broad range of resolutions; however, resolutions finer than 100 bp may slightly reduce prediction performance (Supplementary Table S3).

We further evaluated model performance across different combinations of epigenomic features of histone[24], transcription features [22], and a main transcription factor set consisting of CTCF, RAD21, and SMC3. Models trained on transcription-related feature sets consistently outperformed those trained on histone marks alone, with an improvement of approximately 0.2 in median F1 score, and the combined feature set yielded the highest overall accuracy (Figure 2E, Supplementary Table S4). Transformer models with a smaller number of features for training (e.g., main transcriptional features only, combination of main transcriptional features with histone marks, etc.) also tended to reach an overfitting state more slowly (Supplementary Figure S2). These results are consistent with prior evidence that transcriptional regulators play dominant roles in establishing TAD boundaries. Across all feature combinations at the optimal window length, the Transformer matched or exceeded the performance of the baseline deep learning models, highlighting its ability to integrate positional information of diverse genomic signals (Supplementary Table S4).

**Figure 2.**
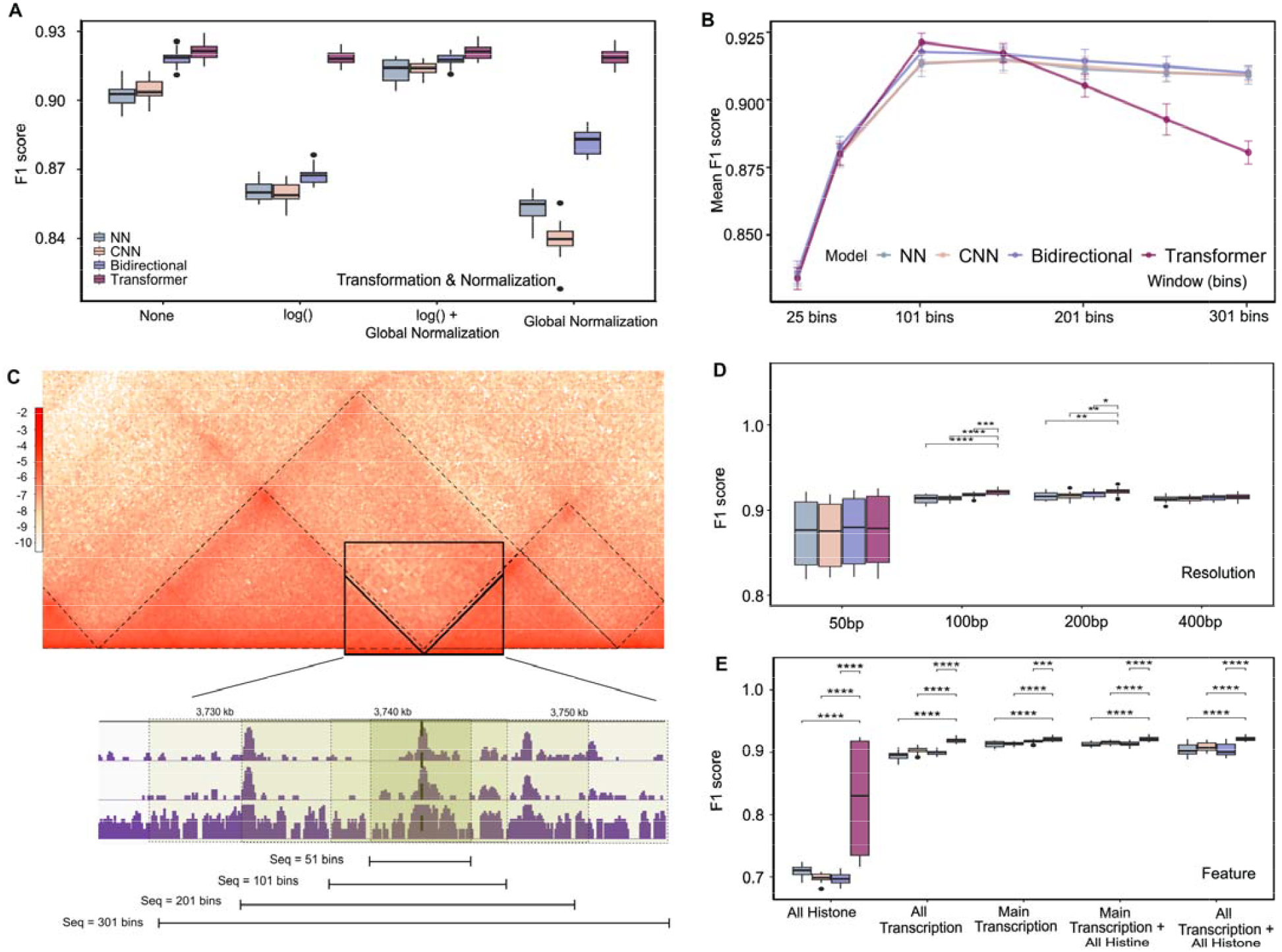
Optimal parameter selection across deep learning architectures (NN, CNN, Bidirectional, Transformer). Model performance assessed by mean and standard deviations of F1 score with other metrics provided in supplementary tables (n = 20 iterations). (A) Normalization effect. (B) Window size (number of bins) effect using the default 100 bp bin size. (C) An example of the GM12878 Hi-C map (zoomed on chromosome 6, 3720kb to 3760kb region), alongside the corresponding 1D genomic transcriptional signal tracks (CTCF, SMC3, RAD21). A representative TAD boundary is highlighted, along with window of increasing sizes capturing additional, and, potentially, irrelevant epigenomic signal. (D) Bin resolution effect using the default 101 bin window size. (E) Epigenomic feature type effect using the default 100 bp bin size and 101 bin windows.

**Supplementary Figure 2.**
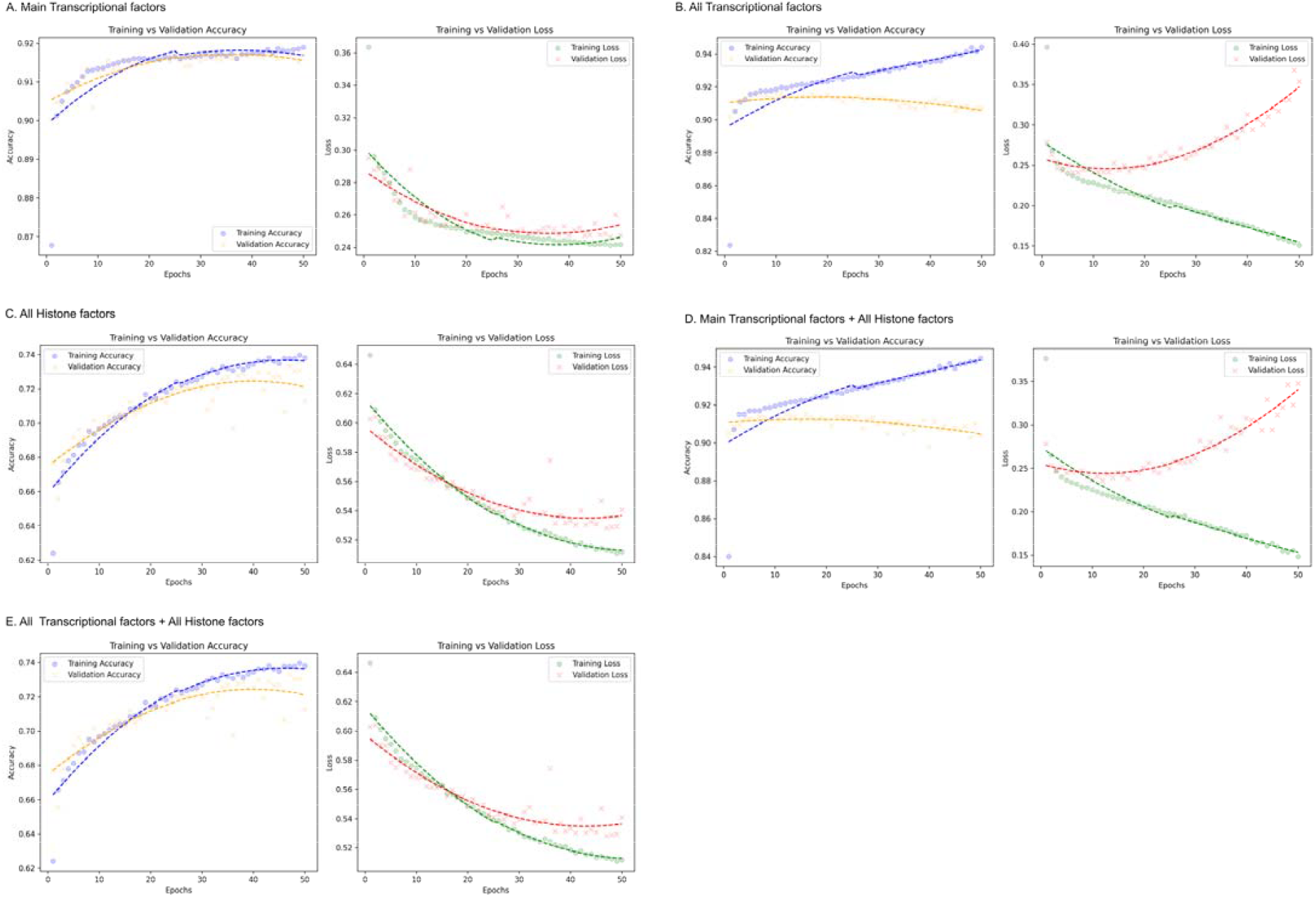
EpiTADformer performance when using various types and numbers of epigenomic features. Diagnostic plots summarize the performance of the Transformer model trained on different feature sets, including main transcriptional factors, all transcriptional factors, all histone factors, and their combinations. For each setting, training and validation accuracy and loss curves are presented.

## Method

### Input Data Representation

Based on our prior studies [25,26], we used genome annotation data from the ENCODE project, also referred to as epigenomic features, for the GM12878 cell line (GRCh38). Transcription factor binding sites (TFBSs) and histone modification marks across 22 chromosomes were used. Any BAM files in the same experiment will be merged by SAMtools (v1.21) [27]. Each chromosome of these merged files was binned into bins of 100 base pairs by default, and the counts of reads in each bin were summarized as total signal values by deepTools (v3.5.6) [28].

TAD boundaries for the GM12878 cell line were identified by preciseTAD, a machine learning method for TAD boundary detection at base-level resolution shown to outperform other methods [22] using CTCF, RAD21, SMC3, and ZNF143. Target bins are defined as bins overlapping with predicted preciseTAD boundary points.

### Definition of Positive and Negative Training Examples

For each target genomic bin, a window of fixed length bins was constructed, where denotes the number of bins included on each side of the target bin at the center of the window (by default, corresponding to 5,000 bp on each side of the center, or a total window length of 10,100 bp). Each window matrix contained rows representing feature total signal values and columns corresponding to bin coordinate positions.

Windows centered on target bins (i.e., bins overlapping preciseTAD boundary points) were defined as positive examples. An equal number of windows centered on randomly selected non-target bins were selected as negative examples. Positive and negative examples were combined and split into training (60%), validation (20%), and testing (20%) sets. Normalization was applied separately to the training, validation, and testing sets, thereby preventing data leakage during model training.

### Normalization

In addition to using raw, unprocessed data, we evaluated three normalization strategies of the input data. Global z-score normalization was applied independently for each chromosome, where the mean and standard deviation were estimated from the chromosome-specific matrix including all genomic regions and the selected features. The natural log transformation transforms the raw signal as *log*(*x* + 0.001) to avoid *log*(0) errors. The combined strategy involves global normalization followed by log transformation.

### Transformer Architecture

We adopted a Transformer-based deep learning model to classify each genomic window as a boundary or non-boundary region. To retain the sequential structure of genomic bins, we added a fixed sinusoidal positional encoding to the window sequence input following the formulation introduced by [29]. For a sequence of *n_bin_* bins and *n_feature_* features, the positional encoding (PE) for a matrix entry at bin position (*pos*) and feature (*f*) was defined as:

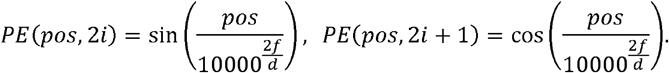

The encoded input, the sum of the original input window matrix and positional encoded window, passed through a Transformer encoder block. Within this block, contextual dependencies are captured through multi-head self-attention, which computes attention weights between all pairs of genomic bins within a window [29]. In our model, we employed two attention heads, each with a head size of 25 for query and key, allowing the network to capture multiple types of regulatory interactions across genomic bins.

The multi-head attention output was combined with the original input through a residual connection and followed by layer normalization. A position-wise feed-forward network (FFN) was then applied, where the first linear layer used a hidden dimension of 250. The output from the FFN was followed by another residual connection and layer normalization. The resulting sequence embedding was aggregated using global average pooling, generating a fixed-length vector (*n*_*features*_ × *n*_*bins*_) summarizing each input window. This embedding was passed through a multi-layer perceptron (MLP) classifier consisting of four dense layers with 650, 450, 164, and 64 units, respectively, each followed by dropout (rate = 0.1). A final sigmoid output node produced the probability that the central target bin is predicted as a boundary (by default, bins with probability larger than the 0.5 threshold are labeled as boundary). At the output layer, to further group detected boundary and non-boundary regions in close proximity, we applied the DBSCAN clustering method. The minimum cluster size (min_sample = 1) was set to one bin, allowing neighboring bins (up to two bins apart, *ϵ* parameter = 200 bp) to be merged into the same cluster, with distances computed in Euclidean space. Clusters thus correspond to genomic regions where bins with similar predictions are dense and nearby.

### Baseline Models

The feed-forward neural network (NN) began with a flatten layer to convert the multi-dimensional input into a 1D vector, which was then processed through two fully connected layers (MLP) with 168 and 100 units, respectively, each activated by ReLU functions. A dropout layer with a rate of 0.3 was incorporated after the first dense layer to prevent overfitting. The final output layer was a sigmoid neuron producing a probability indicating class membership. The model was compiled using the Adam optimizer with a learning rate of 0.0005, employing binary cross-entropy loss.

The convolutional neural network (CNN) was designed for sequential data classification. The model consisted of a single Conv1D layer with 100 filters and a kernel size of 3, followed by a max-pooling layer with a pool size of 1 [30] and a dropout layer (dropout rate = 0.1) to reduce overfitting. The resulting feature maps were flattened and passed through two fully connected (MLP) layers with 224 and 193 hidden units, respectively, both using ReLU activation. The final layer was a single-unit sigmoid neuron that outputs a probability score for binary classification (with a default probability threshold of 0.5). The model was compiled using the Adam optimizer with a learning rate of 0.001 and binary cross-entropy loss.

The bidirectional Long Short-Term Memory (LSTM) network model is tailored for sequence classification. Input sequences were processed through a bidirectional LSTM layer with 100 units, capturing temporal dependencies in both forward and backward directions. The output from the LSTM was fed into an MLP layer of dense layers with 164 and 62 units, each activated by ReLU functions, culminating in a sigmoid output neuron for binary classification. The model was compiled using the Adam optimizer with a learning rate of 0.00035, and the loss function employed was binary cross-entropy.

### Training setup

The models were trained using the Adam optimizer with a fixed learning rate of 0.00005, minimizing the binary cross-entropy loss. To prevent overfitting and ensure stable convergence, we implemented two callback strategies during training. First, a model checkpoint callback was used to save the model weights that achieved the lowest validation loss throughout training. Second, early stopping was applied, monitoring the validation loss with a patience of 10 epochs; training was stopped once the validation loss failed to improve for 10 consecutive epochs, and the model weights were restored to the best-performing checkpoint. The models were then trained for up to 100 epochs (default) using mini-batches of size 17. Various evaluation metrics including accuracy, area under the curve (AUC), precision and recall were collected. All models were implemented in Python (v3.10.18) using TensorFlow (v2.19.0) with the Keras API. Training was performed on 8 Intel / AMD CPU cores with 16 GB RAM of memory.

## Discussion

We developed EpiTADformer, a method to improve TAD boundary detection using epigenomic signal data. By leveraging the concept of region embeddings, we show that epigenomic signals from neighboring genomic bins provides an informative input for deep learning models to detect TAD boundaries. We also show that bin’s positional information can be utilized by the Transformer architecture and enable TAD boundary prediction with high precision.

Despite its strong performance, the Transformer exhibits limitations when the input window becomes too large and covers multiple peak patterns around a boundary. This may happen due to the transformer’s attention to distal patterns associated with other TAD boundaries but not relevant for the target boundary, which is better predicted by local epigenomic signals. In contrast, simpler models may be less affected by window size due to their default prioritization of local patterns and limited memory of distal signals. The Transformer architecture considering proximity of epigenomic signals remains the best performing at the optimal window size (around 10,100 bp).

To address this limitation, future work will explore extended Transformer architectures—for example, whether stacked encoders can improve performance, or whether a multi-input architecture can better resolve short vs. long-range epigenomic patterns. We plan to compare our workflow with more TAD detection baseline methods (e.g., CTCF-RAD21-SMC3 overlapping regions as boundary definition, logistic model, etc.) and recently published methods. We will explore boundary prediction as a binary classification vs. regression problem to establish better thresholds (or lack thereof) for TAD boundary definition. We will test whether a model trained on epigenomic data from one cell line can accurately predict TAD boundaries in another cell line. To further validate our boundary predictions, we will utilize other ground truth datasets (ChIA-PET) to present biological evidence supporting identified boundaries. In summary, our work presents a proof-of-concept that epigenomic signals associated with genomic regions can serve as region embeddings, and their positional relationship can be effectively utilized by the Transformer architecture for TAD boundary predictions.

